# Antibody Binding and Neutralizing Targets within the Predicted Structure of the Poxvirus Multiprotein Entry-Fusion Complex

**DOI:** 10.1101/2025.05.07.652617

**Authors:** Huibin Yu, Wolfgang Resch, Catherine A. Cotter, Wei Xiao, Tase Karamanolis, Ahmed A. Belghith, Maxinne A. Ignacio, Patricia L. Earl, Gary H. Cohen, Bernard Moss

**Affiliations:** Laboratory of Viral Diseases, NIAID, NIH, Bethesda MD, USA; Center for Information Technology, NIH, Bethesda MD, USA; Department of Microbiology, School of Dental Medicine, University of Pennsylvania, Philadelphia, PA, USA

## Abstract

The increased incidence of mpox emphasizes a need for new and improved vaccines. Poxviruses rely on a highly conserved but poorly characterized 11-protein entry-fusion complex (EFC), providing numerous potential targets. Here, we demonstrate that antibodies induced by six of 10 EFC proteins are neutralizing. Protein targets of the neutralizing and non-neutralizing antibodies are located within discrete regions of a model of the EFC predicted by AlphaFold3. Two newly identified targets, A16 and G9, at the apex of the EFC induced cross-neutralizing orthopoxvirus antibodies and protected female mice against a lethal VACV infection. Unexpectedly, antibodies to A16 and G9 were not detected following infection by attenuated or pathogenic VACV, likely due to physical sequestration of the proteins in the viral membrane. Our findings provide a model for the physical, immunogenic and antigenic structure of the EFC, new immunogens for incorporation into recombinant vaccines and suggest a novel poxvirus immune evasion strategy. 150 words.

## Introduction

The recent global outbreak of human mpox, caused by monkeypox virus (MPXV), and the increased incidence of mpox in Africa indicate a need for improved vaccines and therapeutics. The first and second-generation smallpox vaccines are replicating strains of vaccinia virus (VACV) that are contraindicated for those with immunodeficiencies, and have not been used in the current outbreak. The third generation Jynneos vaccine consists of a live, replication-deficient strain of VACV that is safe for a broad range of individuals but is difficult to manufacture at scale, is administered at nearly 1,000 times the dose of replicating strains and requires a booster immunization. Jynneos induces low and non-durable levels of MPXV neutralizing antibody ^1–3^ and provides incomplete protection against mpox ^4–6^. The development of alternative recombinant vaccines requires knowledge of the biology of poxviruses and the identification of optimal immune targets.

Cellular immunity is important in controlling primary orthopoxviral infections, but humoral immunity is sufficient for protection following vaccination ^7–9^. However, the principal targets of the protective antibodies induced by the attenuated viral vaccines remain unknown ^10^. The depletion of only minor amounts of neutralizing antibody from the serum of vaccinees by individual or multiple known antigens led to the hypothesis that a diversity of low-level responses are responsible for protection ^11, 12^. Alternatively, the major neutralizing epitopes may be structurally complex or the major antigens remain to be discovered among the numerous proteins in the virion membrane ^13^. Identification of additional targets of neutralizing antibody could enhance our understanding of poxvirus entry and improve the design of new recombinant vaccines and therapeutics.

The proteins required for entry and membrane fusion of enveloped viruses are the major targets of protective antibodies. Unlike other viruses that utilize one or few viral proteins to enable fusion, poxviruses require 11 highly conserved membrane proteins that form the entry-fusion complex (EFC) in addition to cell-binding proteins ^14^. Comparisons of the EFC protein structures with all proteins in the Protein Database (PDB) and the large database of AlphalFold2 models that cover the complete proteome of humans and other model organisms reveal no similarities, suggesting that the EFC proteins adopt unique folds that are absent or extremely rare in cellular organisms ^15^. Although each of the EFC proteins are potential immune targets, only three are known to induce neutralizing antibodies of which two were shown to be protective in animal models. The present study was initiated to identify targets of neutralizing antibodies that had not been found by analyzing immune serum with protein microarrays ^16, 17^ or by screening large mAb libraries ^18^. By analyzing serum from animals immunized with soluble recombinant EFC proteins, we confirmed the previously identified neutralizing antibodies and discovered three additional ones as well as others that were non-neutralizing. By modeling the 11-subunit EFC with AlphaFold3, we determined that the targets of the neutralizing and non-neutralizing proteins are located in discrete regions. Soluble forms of the two proteins at the predicted EFC apex, A16 and G9, were shown to induce cross-neutralizing antibodies to MPXV and cowpox virus (CPXV) and protect mice against a lethal VACV infection. Despite the ability of A16 and G9 antibodies to neutralize infectious virus, binding antibodies to A16 or G9 were undetected in serum from animals infected with either attenuated or pathogenic VACV. The failure of an immune response to the native proteins is likely due to their physical sequestration within the viral membrane. Our findings provide a model for the physical and antigenic structure of the EFC, new immunogens for incorporation into recombinant vaccines and suggest a novel poxvirus immune evasion strategy.

## Results

### Neutralizing activities of antisera correlate with binding to intact virions

Current information regarding the structure and immunogenicity of the 11 proteins that comprise the VACV EFC is summarized in Table 1. Neutralizing antibodies targeting L1, F9 and A28 were previously reported, and the L1 and A28 antibodies were shown to be protective in animal models. To confirm and extend the analysis, we screened a panel of antisera made by immunizing rabbits with secreted forms of A21, A28, F9, H2, J5, L1 and L5 that were expressed in insect Sf9 cells as described previously for L1 and F9 ^19, 20^. The proteins were purified on Ni-NTA beads, mixed with Freund’s adjuvant and injected at subcutaneous and intramuscular sites of rabbits and boosted at 2-week intervals (Fig. 1a). In each case, the presence of specific antibodies was confirmed by binding to proteins of the predicted size on SDS-polyacrylamide gel (PAGE) blots of lysates from VACV-infected cells (Fig. 1b) and by a quantitative enzyme-linked immunosorbent assay (ELISA) to dissociated purified virions (Fig. 1c). A flow cytometry assay was used to determine the titers of antibodies that neutralized the infectivity of VACV expressing green fluorescent protein (GFP) ^21^. Anti-L1, –F9, –A28, and –J5 sera neutralized VACV, but no neutralization above the control was found for the anti-H2, –L5 or –A21 sera (Fig. 1d), even though the latter sera bound well to dissociated virus particles. The ability of the antisera to bind intact virions was also determined. To account for the differences in titers of the different sera, the data were normalized using the values for binding to dissociated virions. The anti-A28, –F9, –L1 and –J5 sera exhibited ∼100-fold greater binding to intact virions than the anti-H2, –L5 and –A21 sera (Fig. 1e), similar to the differences in neutralizing activities. Thus neutralization and binding to intact virions were correlated and of the seven antisera analyzed, four were neutralizing and three were non-neutralizing.

**Fig. 1.**
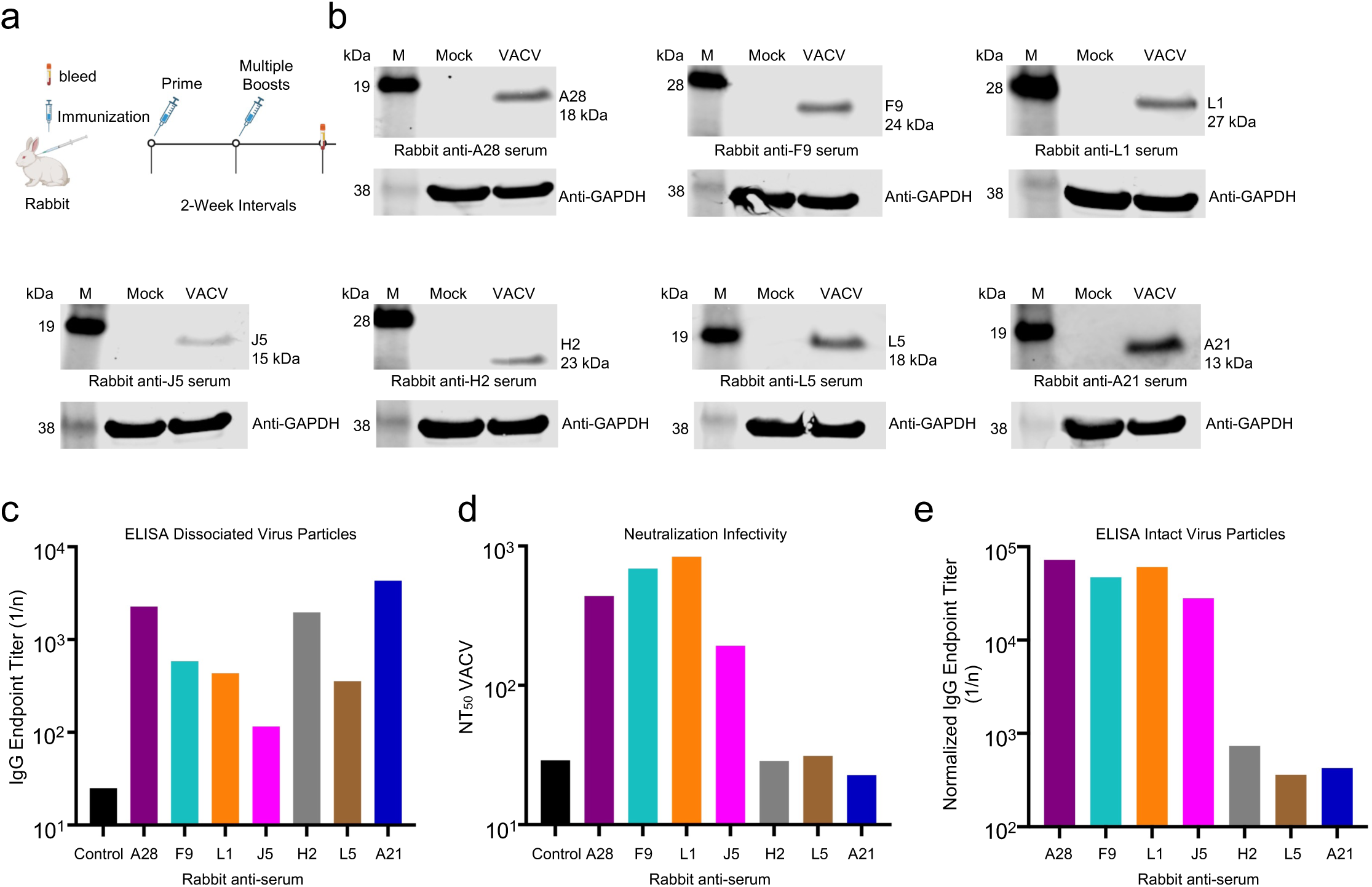
Binding and neutralizing activities of antibodies to EFC proteins. (**a**) Antisera from rabbits that were immunized multiple times at 2-week intervals with individual purified EFC proteins were used for immunoasssays. (**b**) SDS-PAGE blots of lysates of Mock-or VACV-infected cells were probed with rabbit antisera. EFC protein masses are indicated on the right. M, refers to marker proteins with masses on left. GAPDH was detected as a loading control. **(c)** Binding of rabbit antisera to purified VACV particles that were dissociated with SDS, renatured, and adsorbed to wells of an ELISA plate. **(d)** Neutralization of VACV by rabbit antisera. **(e)** Binding of rabbit antisera to purified intact VACV particles. Data were normalized using the binding titers for dissociated virions. ELISA and neutralization titers determined in duplicate.

**TABLE 1.** EFC proteins.

| VACV name | OPG number | kDa | TM<sup>a</sup> | PDB ID | Neut Ab | Paralog | Ref |
| --- | --- | --- | --- | --- | --- | --- | --- |
| A16 | 143 | 43 | C | 8GP6 (A16/G9)<sup>b</sup> | ? | G9, J5 | 42, 52 |
| A21 | 147 | 14 | N | 8UOR | ? |  | 38 |
| A28 | 155 | 16 | N | 8GQO | + |  | 53, 54 |
| F9 | 053 | 24 | C | 6CJ6 | + | L1 | 20, 55 |
| G3 | 086 | 13 | N | 7YTT (G3/L5)<sup>b</sup> | ? |  | 56, 57 |
| G9 | 094 | 39 | C | 8GP6 | ? | A16, J5 | 39, 52 |
| H2 | 107 | 22 | N | 8INI | - |  | 58, 59 |
| J5 | 104 | 15 | C | 8WT5 | ? | A16, G9 | 60 |
| L1 | 095 | 27 | C | 1YPY | + | F9 | 61, 62 |
| L5 | 099 | 15 | C | 7YTT (G3/L5)<sup>b</sup> | ? |  | 37, 57 |
| O3 | 076 | 4 | N | - | ? |  | 63 |

<sup>a</sup> N- or C-terminal transmembrane domain; <sup>b</sup> Structure determined for heterodimer

### A16 and G9 are targets of neutralizing antibodies

To determine whether additional EFC proteins are targets of neutralizing antibodies, we expressed secreted forms in mammalian cells as outlined for VACV A16 and G9 (Fig. 2a). Transmembrane (TM) domains were deleted and signal peptides and epitope tags added. Predicted N-glycosylation sites were mutated because A16 and G9 do not normally traffic through the secretory pathway. As A16 and G9 form a stable heterodimer, the open reading frames (ORFs) were inserted individually or together in a dual promoter expression plasmid (Fig. 2b). Following transfection of plasmids into human 293T cells, expression of A16-Avi-His and G9-Flag were detected by SDS-PAGE immunoblots of cell lysates and concentrated media (Fig. 2c). A16 was mostly cell-associated when expressed alone, but secretion increased when co-expressed with G9 suggesting an intracellular interaction between the two proteins. In contrast, G9 was secreted to a similar extent when expressed alone or together with A16 (Fig. 2c). Thus G9 appears to act as a chaperone for A16.

**Fig. 2.**
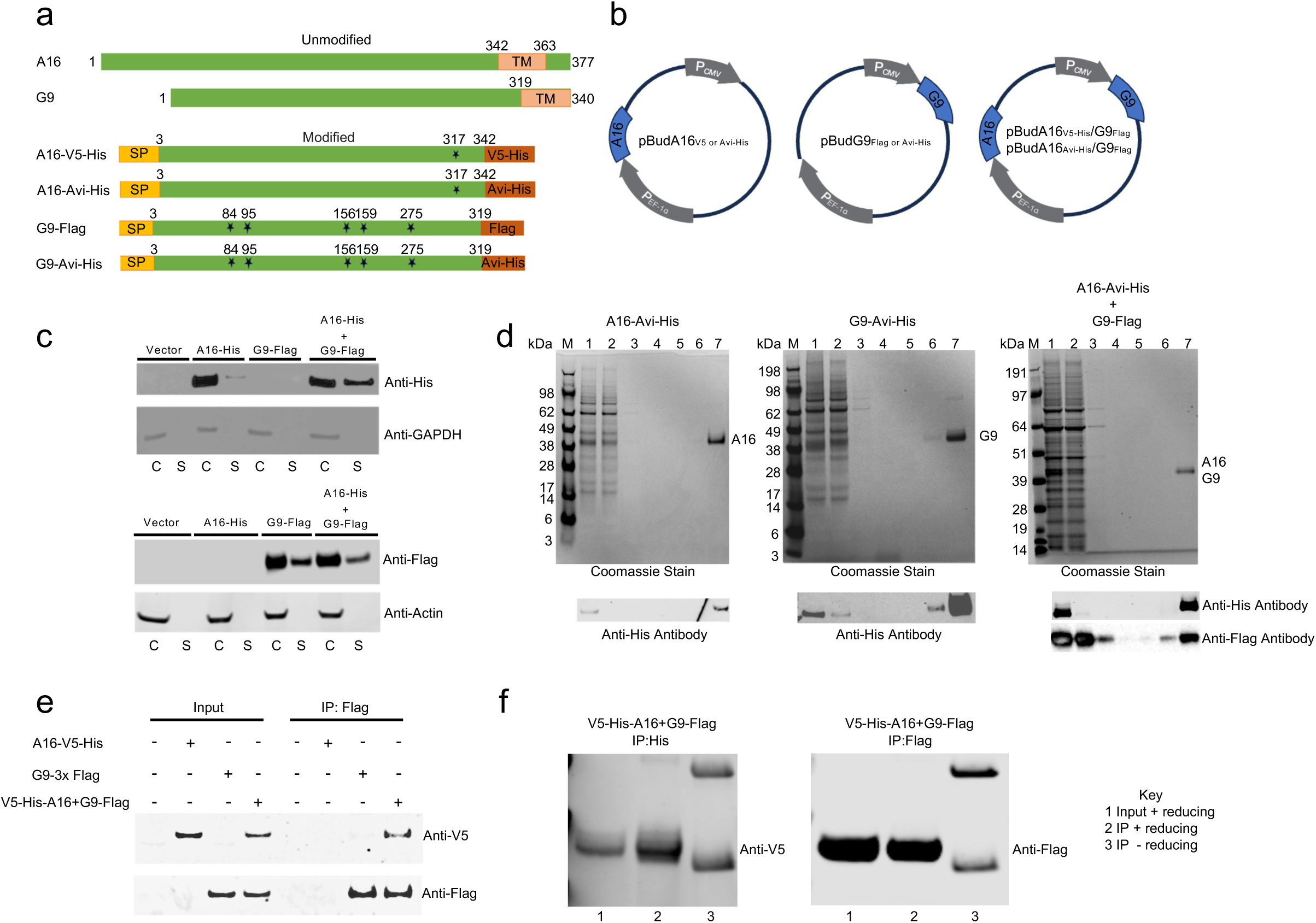
Expression and purification of A16 and G9 proteins. **(a)** Unmodified full-length A16 and G9 proteins (top) and modified proteins (bottom) showing removal of the TM domain, addition of a signal peptide (SP), potential N-glycosylation site of A16 (asterisks) changed from threonine to valine and of G9 changed serines and threonines to alanines, and C-terminal epitope tags. **(b)** pBudCE4.1 vector containing modified A16, G9, and combined A16/G9 ORFs regulated by CMV or EF-1α promoters. **(c)** Immunoblots of cell lysates (C) and secreted proteins (S) following transfection of plasmids into Expi293F^TM^ cells. Primary antibodies used to probe blots indicated on right. **(d)** Purification of proteins on Ni-NTA beads. SDS-PAGE of starting material (1), flow through (2), wash (3-5) and elution (6, 7) fractions detected by Coomassie blue staining and with anti-His and anti-Flag antibodies. Marker proteins (M). (**e)** Immunoblot of SDS gel probed with anti-V5 and anti-Flag antibodies showing co-purification of secreted A16 and G9 with anti-Flag agarose beads from 293T cells transfected with plasmids expressing A16-V5-His, G9-3XFlag, and A16-V5-His+G9-3XFlag. **(f)** Detection of secreted A16/G9 dimer from 293T cells transfected with plasmid expressing A16-V5-His+G9-3XFlag after SDS treatment in presence or absence of reducing agent. Representatives of two or more immunoblots shown.

Samples containing A16-Avi-His, G9-Avi-His or A16-Avi-His + G9-Flag were purified by affinity chromatography on Ni-NTA beads and analyzed by SDS-PAGE (Fig. 2d). Coomassie blue stained bands corresponding to the expected masses of the tagged proteins were detected. In the case of A16-Avi-His + G9-Flag, both proteins were bound to and eluted from Ni-NTA beads even though only A16 had a His tag indicating that they formed a heterodimer. Immunoblotting was needed to identify A16 and G9 when analyzed together because of their similar mass and electrophoretic mobilities when denatured with SDS and a reducing agent (Fig. 2d). Additionally, A16-V5 was pulled down with anti-Flag when co-expressed with G9-Flag confirming that they formed a heterodimer (Fig. 2e). In the experiments described thus far, the proteins were denatured with SDS and a reducing agent before SDS-PAGE. In the absence of reducing agent, some of the co-expressed A16 and G9 migrated as a heterodimer (Fig. 2f), presumably due to numerous intramolecular disulfide bonds that prevented complete denaturation by SDS alone.

There were no previous reports regarding the immunogenicity of A16 and G9 and for this reason we analyzed the induced antibodies in detail. Female BALB/c mice were injected subcutaneously (SC) with purified A16, G9 or A16/G9 heterodimer plus AddaVax adjuvant (Fig. 3a). Mice were also immunized with MVA, the attenuated strain of VACV used as smallpox and mpox vaccines. Antibodies in the serum of mice receiving A16, G9 or A16/G9 bound to the corresponding antigen and A16/G9, which increased after successive boosts (Fig. 3b, S1a). By contrast, no A16 or G9 binding antibodies were detected in serum of mice immunized with MVA suggesting that A16 and G9 have low immunogenicity in the context of viral infection, as will be confirmed and extended in a later section.

**Fig. 3.**
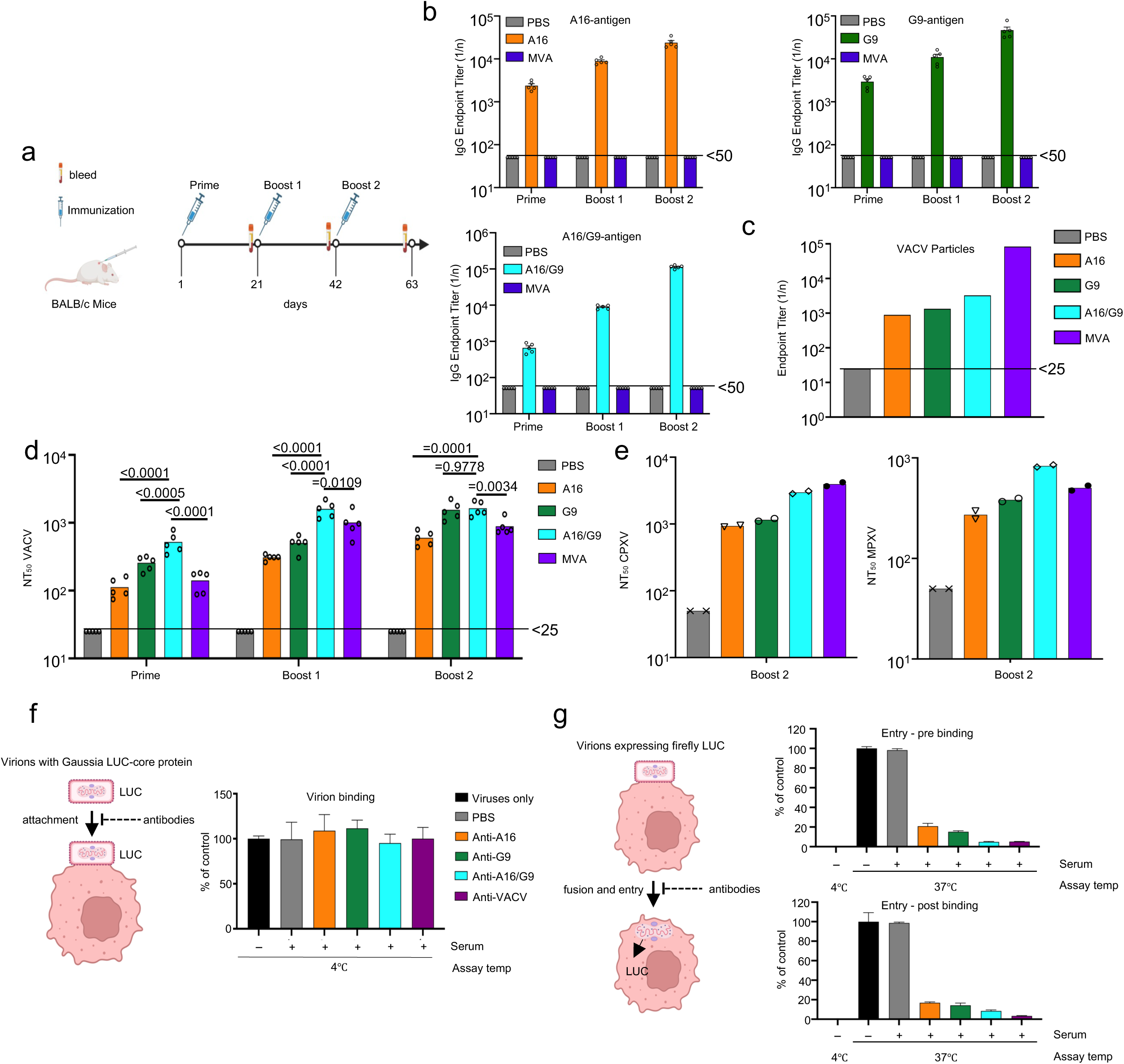
Binding and cross-neutralizing activities of A16/G9 antibodies. **(a)** Groups of mice (n=5) received three 10 μg doses of A16, G9, or A16/G9 heterodimer SC with AddaVax or 10^7^ PFU of MVA IM. **(b)** ELISA assessing binding of immune sera from individual mice to A16, G9 and A16/G9 proteins. **(c)** ELISA assessing binding of pooled immune sera to purified intact VACV. **(d)** VACV neutralization titers determined with sera from individual mice. Significance (p values) calculated by one-way ANOVA with multiple comparisons test. **(e)** CPXV and MPXV NT50 determined in duplicate on pooled sera. **(f)** Effects of mouse antisera on binding to cells of Gaussia luciferase recombinant VACV. Experiment was carried out in triplicate. Error bars are SEM. **(g)** Effects of mouse immune sera added before or after attachment of WRvFire to cells and entry assessed by luciferase activity. Experiment was carried out in triplicate. Error bars are SEM.

Antisera to A16, G9, A16/G9 and MVA bound to purified intact virions as determined by ELISA with anti-MVA sera having the highest binding presumably because of antibodies to numerous proteins (Fig. 3c). The anti-A16/G9 serum exhibited significantly higher neutralizing activity than that of A16 and G9 until the second boost when G9 caught up (Fig. 3d). Despite the relatively high amount of virion-binding antibody produced by MVA, the neutralization titer was significantly less than for A16/G9. The VACV A16, G9 and A16/G9 antisera cross-neutralized MPXV and CPXV (Fig. 3e) consistent with the high sequence conservation of these entry proteins.

Depletion experiments demonstrated that the anti-A16/G9 sera contained antibodies that bind both A16 and G9 antigens. Magnetic beads coated with A16 protein or G9-protein completely removed antibodies that bind to A16 or G9 antigen, respectively, whereas A16/G9 coated beads removed both (Fig. S1b,c). Moreover, only beads coated with A16/G9 completely removed antibodies that bind to A16/G9 antigen (Fig. S1d). Similar results were obtained when neutralization was analyzed: only beads coated with A16/G9 completely removed neutralizing activity (Fig. S1e). Thus, the heterodimer induces antibodies to both A16 and G9.

Further studies were carried out to analyze the mode of virus neutralization. To determine whether antibodies inhibit cell binding, immune serum was incubated with a recombinant VACV in which the A4 core protein is fused to Gaussia luciferase ^22^. After allowing the virus to adsorb to cells in the cold, unattached virus was removed and luciferase activity determined. None of the immune sera reduced virus binding to levels appreciably lower than control serum (Fig. 3f).

An entry assay was based on the packaging of a complete transcription system in infectious VACV particles that enables expression of early proteins after core entry. Sera from vaccinated mice were incubated either prior to or after cell binding of the recombinant virus WRvFire, which expresses luciferase from an early promoter. Either way, antiserum to A16, G9, A16/G9, and infectious VACV severely inhibited luciferase activity (Fig. 3g). We concluded that anti-A16 and anti-G9 antibodies can prevent virus entry at pre-or post-binding steps.

### MPXV homologs of VACV G3 and L5 are non-neutralizing

Rabbit antiserum to VACV L5 was non-neutralizing and bound poorly to intact virions (Fig. 1d, e). However, antibody to G3 a binding partner of L5 ^23^ was not tested and it is possible that the complex is more immunogenic than either protein alone as is true for the H2-A28 heterodimer ^24^. For reasons unrelated to this study, we expressed MPXV M5 and G2 which differ from their VACV L5 and G3 homologs by only 1 and 2 amino acids, respectively. Like with A16 and G9, the proteins were modified by removing the TM domain, adding a signal peptide and epitope tags, and mutating predicted N-glycosylation sites (Fig. 4a). Modified G2 and M5 ORFs were inserted into a dual promoter plasmid, which was transfected into human 293T cells (Fig. 4b).

**Fig. 4.**
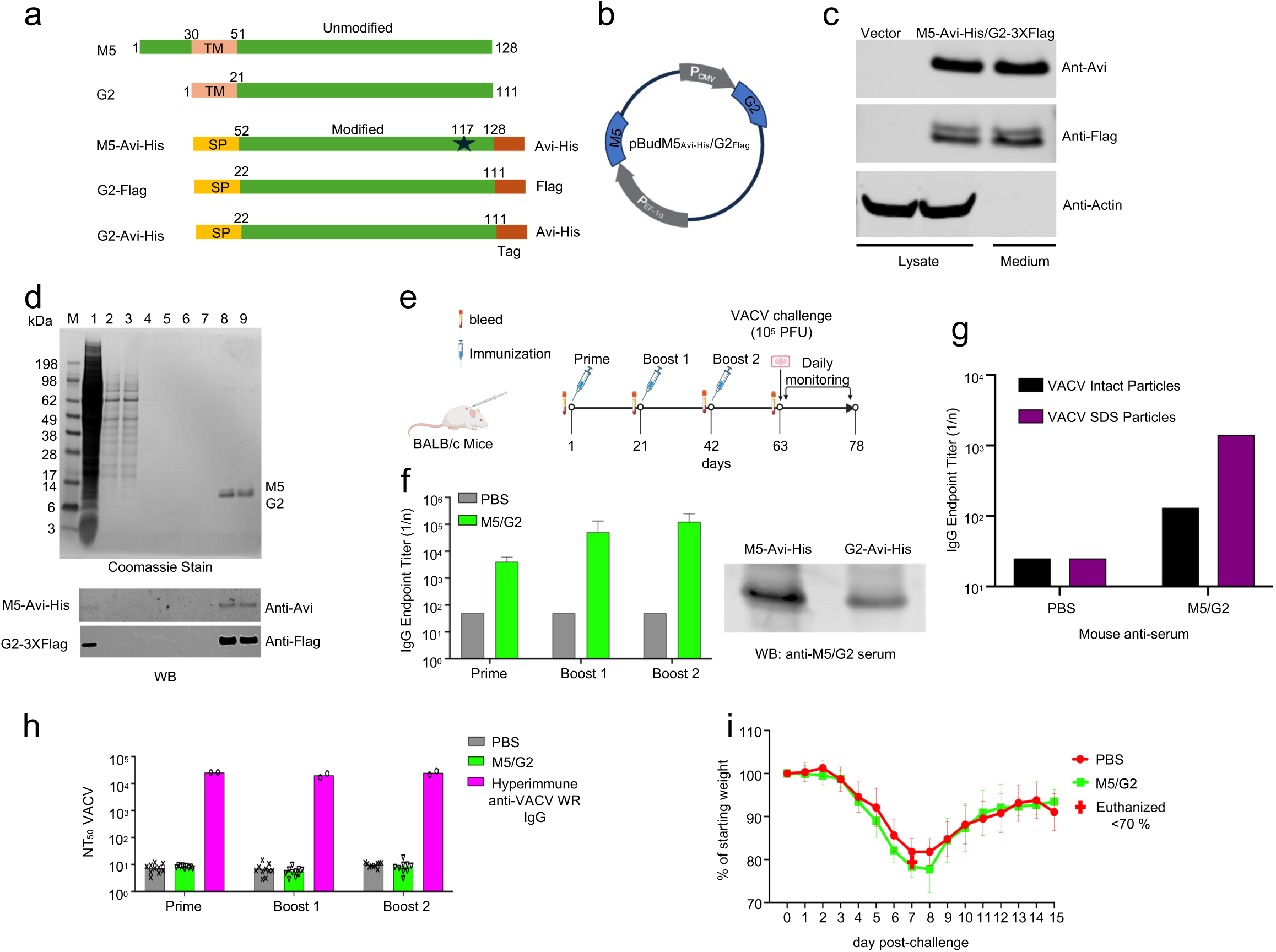
Immunogenicity of MPXV M5 and G2 proteins. **(a)** Representation of unmodified full-length MPXV M5 and G2 proteins and modified proteins. Asterisk indicates serine to alanine mutation of N-glycosylation site. **(b)** Diagram of pBudCE4.1 expression vector with M5 and G2 ORFs. **(c)** Immunoblot of lysate and medium of Expi293F^TM^ cells transfected with M5-Avi-His/G2-3XFlag plasmid. Primary antibodies used to probe blots shown on right. **(d)** SDS-PAGE analysis of proteins at steps during Ni-NTA affinity purification of secreted M5/G2 heterodimer. Lanes: (1) cell lysate, (2) medium, (3) flow through, (4-7) wash, (8, 9) eluate. Coomasie blue stain on top, immunoblots below. **(e)** Experimental plan for immunization with M5/G2 and VACV challenge with 10^5^ PFU of VACV-WR. **(f)** ELISA demonstrating binding to M5/G2 of serum antibody after prime and boosts. SDS-PAGE blots of purified M5-Avis-His and G2-Avi-His probed anti-M5/G2 serum. **(g)** Binding of anti-M5/G2 serum to intact and SDS-disrupted VACV particles. **(h)** Absence of VACV neutralizing antibody in serum from M5/G2 immunized mice. IgG from VACV immunized rabbit was used as a positive control. (i) Weight loss plotted as percent of starting weight of mice (n=5) following challenge with 10^5^ PFU of VACV. Bars represent SD.

Expression of M5-Avi-His and G2-Flag were detected by probing SDS-PAGE immunoblots of cell lysates and concentrated media with antisera to the epitope tags (Fig. 4c). The heterodimer was purified on Ni-NTA beads (Fig. 4d), mixed with AddaVax adjuvant and used to immunize mice (Fig. 4e). Antibdies to the heterodimer measured by ELISA were detected after the prime immunization and increased after boosting and a Western blot verified that the anti-M5/G2 serum bound to both M5 and G2 (Fig. 4f). Binding values to dissociated virus particles was 11-fold higher than to intact virus particles indicating low accessibility of antibodies to the latter (Fig. 4g) and no VACV neutralizing activity was detected even after a prime and two boosts (Fig. 4h). In addition, mice immunized with the heterodimer lost weight similar to control mice following an intranasal (IN) challenge with 10^5^ PFU of VACV (Fig. 4i). We concluded that M5 and G2, and likely their close VACV homologs L5 and G3, are non-neutralizing and non-protective.

### In-vitro neutralizing antibodies to G9 and A16 correlate with protection in an animal model

A major objective of this study was to define previously unrecognized immune targets that could be used in a new generation of recombinant orthopoxvirus vaccines. The strong neutralizing antibody response to the A16 and G9 proteins made them good candidates. However, previous studies with the A27 virion-binding protein had shown that high *in vitro* neutralization of VACV may not correlate with protection *in vivo* ^25, 26^. Therefore, following priming and boosting with A16, G9 and A16/G9 (Fig. 5a), the mice were challenged IN with 10^5^ PFU of VACV. We used WRvFire, a derivative of VACV WR that retains pathogenicity and expresses firefly luciferase to allow bioluminescent imaging (BLI) of virus replication and spread ^27, 28^. The PBS control group exhibited severe weight loss during the first 8 days with three of ten exceeding 30% and requiring euthanasia (Fig. 5b). All immunized mice survived with the each group losing significantly less weight than the PBS control (p<0.0001) as determined for the combined days 7 to 9 by one-way ANOVA with Tukey’s multiple comparison test. In addition, the A16/G9 group lost significantly less weight than the A16 or G9 groups (p<0.0001), whereas there was no significant difference between the A16 and G9 groups (p=0.9899).

**Fig. 5.**
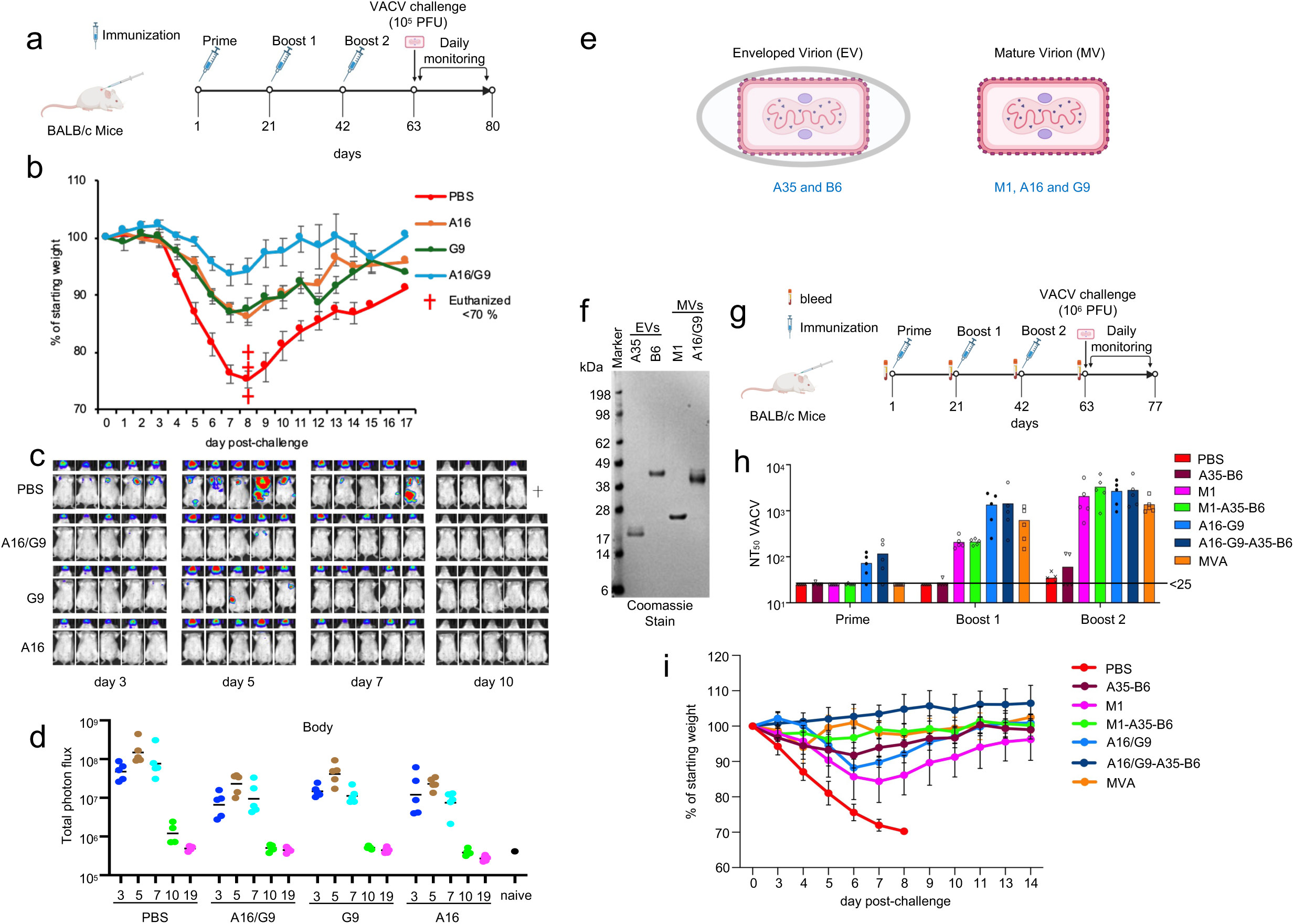
Protection of A16– and G9-immunized mice against VACV infection. **(a)** Groups of female mice from two independent experiments (n=10) were primed and boosted twice with 10 µg of A16, G9 or A16/G9 proteins or with 10^7^ PFU of MVA and challenged IN with 10^5^ PFU of VACV-WR. **(b)** Percent of starting weight post-challenge. † indicates death. Bars are SD. **(c)** BLI of heads and bodies of immunized mice post-challenge with WRvFire from one of two similar experiment (n=5). **(d)** Photon flux of bodies of mice imaged in panel c. Bars represent mean. **(e)** Diagram of orthopoxvirus EV and MV indicating MPXV A35 and B6 proteins of EV outer membrane and MPXV M1 and VACV A16 and G9 proteins of MV membrane. **(f)** Coomassie blue stained SDS gel showing purity of recombinant proteins used for immunization. kDa of marker proteins shown at left. **(g)** Groups of mice (n=5) were primed and boosted twice with 10 µg of recombinant MV and EV proteins or 10^7^ PFU of MVA. **(h)** Anti-VACV neutralizing titers of serum from individual mice after prime and boosts. LOD of 25 indicated. **(i)** Percent of starting weights following challenge with 10^6^ PFU of VACV. Immunogens indicated on right. Bars represent SD.

To evaluate virus replication and spread, luciferin was injected intraperitoneally, and BLI visualized. As luciferase has a half-life of < 2 h, BLI measures virus infection near the time of analysis. Because of variations in light quenching by different tissues, BLI of head and body cannot be directly compared to each other and split images of mice are shown (Fig. 5c). The control PBS mice exhibited BLI in the heads and chest on day 3 that increased and spread to the abdomen on day 5 and declined by day 10 in surviving mice. Mice immunized with A16, G9 or A16/G9 had BLI in the heads that diminished earlier than in the controls and few had discernible BLI in the chest or body. Quantitative photon flux measurements confirmed significantly lower (p<0.016) luminescence in the bodies on each day for mice immunized with A16/G9 and G9 and for all except day 3 for mice immunized with A16 compared to the PBS controls as determined by one-way ANOVA with multiple comparisons. Thus, immunization with A16/G9 prevents virus spread but not initial replication at the IN site of infection.

A16 and G9 and other EFC proteins are components of the mature virion (MV). Orthopoxviruses have a second infectious form called the enveloped virion or EV with an additional outer membrane that must eventually be disrupted to allow the EFC to mediate fusion with cell membranes (Fig. 5e). EVs are important for virus spread as well as for shielding the internal MV from neutralizing antibodies. Previous studies showed that antibodies to an MV protein such as L1 can provide partial protection, but that more complete protection requires antibodies to EV proteins as well ^19^. Because of availability, we used purified MPXV A35 and B6 EV proteins with 7 and 10 amino acid differences from the VACV homologs A33 and B5 (Fig. 5e, f). For comparison with A16/G9, we used purified MPXV M1 MV protein, which has 2 conservative amino acid differences from the VACV L1 protein and has been used in mRNA and protein VLP vaccines (Fig. 5e, f). Mice were vaccinated with combinations of MV and EV proteins and with MVA to provide a positive reference control (Fig. 5g). A16/G9 induced higher neutralizing antibodies than M1 after the prime and first boost although the latter caught up at the second boost (Fig. 5h). As expected the EV proteins did not induce MV-neutralizing antibodies alone or enhance their formation when combined with A16/G9. When the mice were challenged IN with 10^6^ PFU of VACV-WR, the control mice rapidly lost weight and succumbed whereas all immunized mice survived with varying weight loss (Fig. 5i). Mice immunized with A16/G9 alone lost significantly less weight than M1 alone (p=0.0368) when combined days 6 to 8 were analyzed by one-way ANOVA with Tukey’s multiple comparison test. However A16/G9 plus A35 and B6 provided similar protection as M1 + A35 and B6 (p=0.997) and MVA (p=0.9717). Thus, A16/G9 provided better protection than M1 and synergized with EV proteins supporting potential use in a multivalent vaccine.

### Antibodies to A16 and G9 are not elicited during VACV infection

In the experiment of Fig. 3b, antibodies binding to A16 or G9 were not detected in mice immunized with MVA. To extend this observation, we analyzed sera from mice and rabbits that were immunized with the pathogenic replicating WR strain of VACV as well as from non-human primates immunized with replicating (ACAM2000) or non-replicating (MVA-BN) vaccine strains intradermally or subcutaneously. In none of these sera were anti-A16 or –G9 binding antibodies detected (Fig. 6a). Nevertheless, pathogenic and vaccine strains of VACV produced antibodies to other VACV or MPXV EFC proteins (A28, M1, F9) and to the H3 virion attachment protein.

**Fig. 6.**
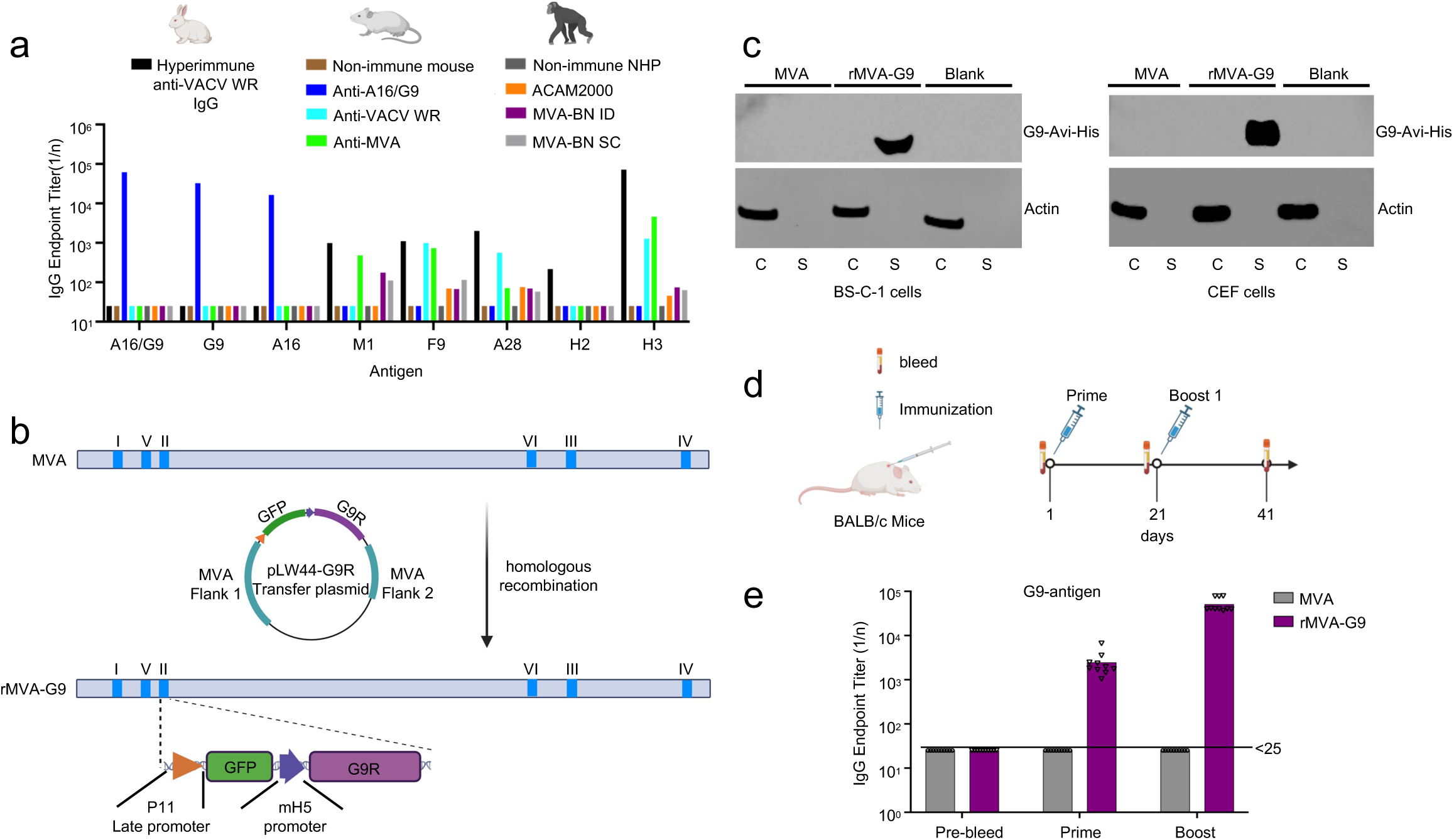
Deficient antibody response to A16 and G9 by VACV. **(a)** Binding of sera from VACV-infected animals to VACV G9, A16, F9, A28, H2, and MPXV M1 and H3 (homologs of VACV L1 and H3, respectively) proteins. The following pooled serum samples were tested: hyperimmune IgG from rabbits infected multiple times with VACV-WR; nonimmunized mice or mice inoculated percutaneously with 10^5^ PFU of VACV-WR, or primed and boosted twice with 10^7^ PFU MVA IM or 10 µg A16/G9 with AddaVax SC, unimmunized rhesus macaques or macaques immunized with 10^6^ PFU of ACAM2000 percutaneously or 10^8^ PFU of MVA-BN ID or SC. **(b)** Construction of recombinant MVA (rMVA-G9) by homologous recombination. The six deletion sites in MVA genome indicated by Roman numerals. The secreted form of G9-Avi-His DNA (Fig. 2a) was inserted into the LW44 shuttle vector under control of the mH5 early/late promoter, which allowed recombination into deletion II, and the GFP-expressing virus was clonally purified. **(c)** Immunoblot analysis of G9-Avi-His in the cell lysates and medium of BS-C-1 and CEF cells infected with rMVA-G9. **(d)** Immunization of BALB/c mice (n=10) with 10^7^ PFU of parental MVA and rMVA-G9. **(e)** Analysis of anti-G9 antibodies by ELISA in the serum of individual immunized mice (n=10).

We considered two general mechanisms for the failure of live VACV to induce antibodies to A16 and G9: active immune suppression or passive concealment. To investigate these possibilities, a secreted form of G9 was inserted into MVA by homologous recombination (Fig. 6b). The native G9 lacks a signal peptide and is inserted into the viral membrane within the cytoplasm, whereas the addition of a signal peptide and removal of the TM domain should drive the recombinant G9 through the secretory pathway. G9 was efficiently secreted into the medium following infection of permissive chicken embryo fibroblast (CEF) or non-permissive African green monkey kidney BS-C-1 cells (Fig. 6c). Expression of the secreted G9 had no effect on MVA plaque formation in CEFs, and viral replication in either CEF or BS-C-1 cells (Fig. S2a,b). Antibodies binding to G9 were detected in the serum of mice infected with MVA-G9 after a single vaccination and boosted by a second, whereas none was detected in the serum of mice infected with parental MVA (Fig. 6e). These data are consistent with MVA preventing antigen presentation by concealing or sequestering A16/G9 when presented in the viral membrane-associated EFC, although other explanations are not excluded.

### Location of targets of neutralizing and non-neutralizing antibodies on a model of the EFC

Although structures for the ectodomains of all EFC proteins except for the 35 amino acid O3 have been determined (Table 1), there is only limited information regarding their organization, as the EFC has not yet been isolated in a native form nor can it be recognized as a discrete structure on the surface of virions. Nine of the EFC proteins (A16, A21, A28, G3, G9, H2, J5, L5, O3) are considered core components as the absence of expression of any one prevented assembly of the EFC. Assuming one copy of each protein, a complex of the nine core proteins (Fig. 7a) and a holocomplex of all 11 proteins (Fig. 7b) were predicted with AlphaFold3 ^29^. An overlay of the colored holocomplex on the uncolored core complex shows some minor differences in the predicted structure caused by inclusion of L1 and F9 (Fig. 7c). Several methods were used to evaluate confidence. The predicted local distance difference test (pLDDT) is a metric used by AlphaFold to estimate local confidence at the level of individual amino acid residues increasing from 0 to 100. The predicted aligned error (PAE) is effectively a measure of how confident AlphaFold is that the domains are well packed and that the relative placement of the domains and subunits are correct. Both metrics provided high confidence overall (Fig. S3), although the placement of L1 relative to the other subunits has a higher predicted error. The available ectodomain structures from the Protein Database (PDB) could be superimposed on the holocomplex (Fig. S4a-d); the root mean square deviation (RMSD) of the pruned structural alignments ranged from 0.473 to 1.07 Å (Table S1) indicating good correspondence. In addition, 25 of the 34 known disulfide-bonded sulfur atom pairs were within 2.05 Å of each other and 33 were within 2.15 Å in the model, close enough to form disulfide bonds (Fig. S4e). Furthermore, the predicted adjacency of external domains of A16 and G9, A28 and H2, and L5 and G3 in the model is consistent with experimental data demonstrating that these pairs form stable heterodimers ^23, 30, 31^. Novel features of the model include the clustering of all TM domains and the alignment of parallel β-sheets of A16, G9, J5, F9 and L1 to form a hinge-like structure (Fig. S5) that could modulate structural changes. The Fig. S1 movie shows a 3-dimensional space-filling model of the holocomplex, surface lipophilicity, and the parallel β-sheets.

**Fig. 7.**
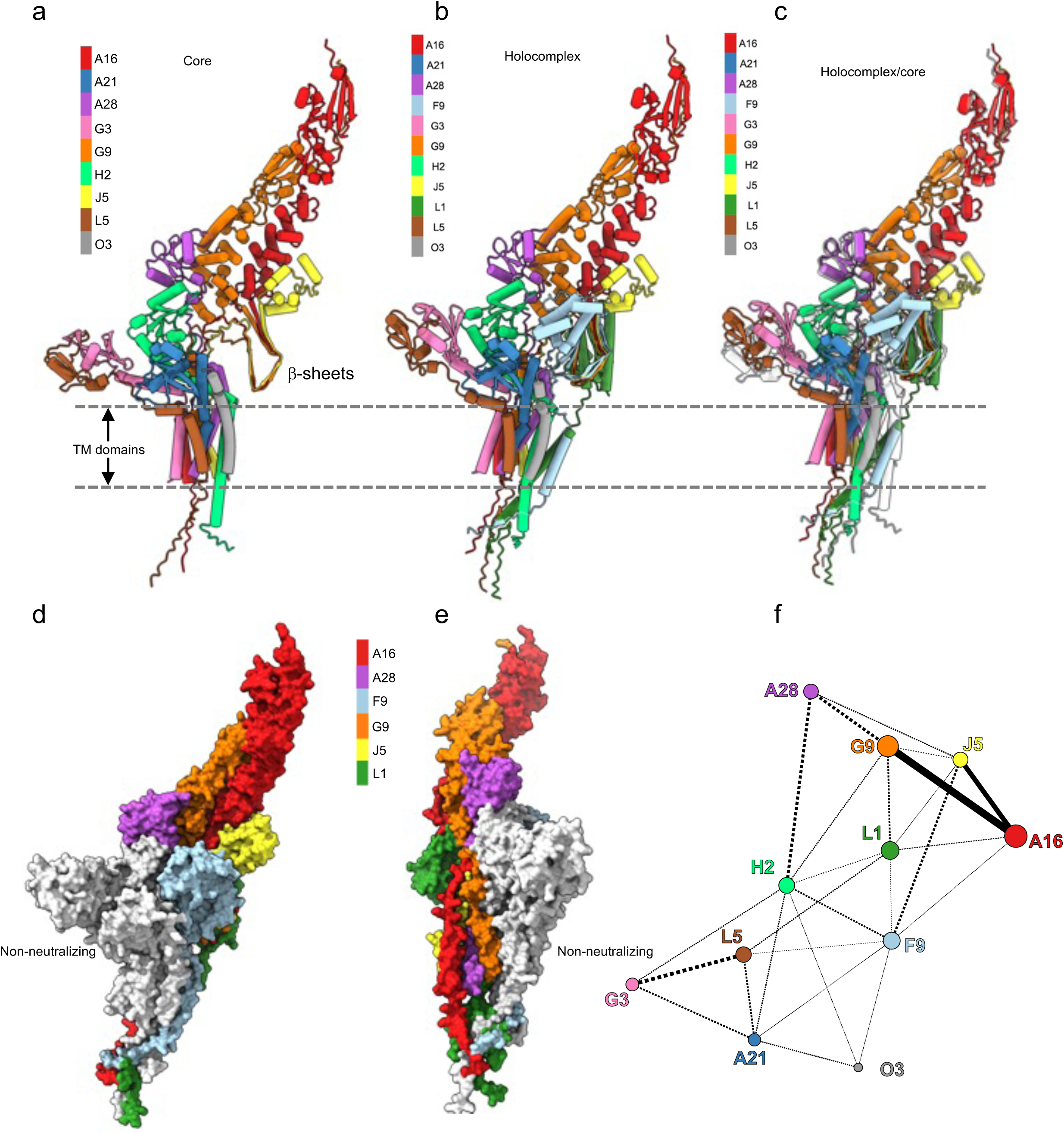
Antibody targets on EFC model prediceted by AlphaFold3. Ribbon structural models of the VACV EFC 9-protein core **(a)**, 11-protein holocomplex **(b)**, and holocomplex superimposed on uncolored core **(c)** predicted by AlphaFold3. Dashed lines indicate locations of TM domains. Individual EFC proteins identified in color keys. Space filling models shown in two orientations **(d, e)** of EFC holocomplex with neutralizing targets in color and non-neutralizing targets in gray. Individual neutralizing targets identified in color key. **(f)** Diagram of buried solvent-accessible surface area (SASA) between pairs of proteins. The line widths were manually scaled to the size of the SASA. Dotted lines are less than half of the maximum SASA value.

Next we mapped the regions of the EFC that are targets of neutralizing and non-neutralizing antibodies. The proteins L1, F9, A28, A16, G9 and J5 that are targets of neutralizing antibodies are shown in color, whereas the non-neutralizing region comprising proteins L5, A21, G3, H2 is depicted in gray in two dimensions (Fig. 7d,e) and in three dimensions (S2 movie). A16 and G9, which induce strongly neutralizing antibodies form the apex of the EFC with J5, A28, F9 and L1 below. Associations between the neutralizing target proteins and between the non-neutralizing target proteins is supported by the network diagram showing the buried solvent accessible surface areas (SASA) (Fig. 7f). The strongest predicted interactions are between G9, A16 and J5, although the latter has not been determined experimentally. An interaction between A28 and G9 is predicted in addition to the known interaction of A28 and H2. The non-neutralizing protein targets form a contiguous region closer to the base of the EFC in which G3 and L5 are known to interact and are predicted to interact with A21. The correlation of antibody neutralization with antibody binding to intact virus particles suggests that the non-binding region may be buried or masked by other proteins.

## Discussion

This study was initiated with three objectives: identify novel EFC targets of neutralizing and protective antibodies, predict the organization of the EFC using AlphaFold, and localize the neutralization sites within the EFC model. In addition to meeting these objectives, an unanticipated finding was that antibodies to at least two components of the EFC, A16 and G9, are not detected after infection with live VACV even though neutralizing antibodies to these proteins were induced by the soluble proteins.

The previously reported immunogenicity of individual EFC proteins is listed in Table 1. L1 ^32–34^, A28 ^35^ and F9 ^20^ were previously shown to be targets of neutralizing antibodies, though antibody to the latter was not evaluated for protection. Of these only A28 is an EFC core component. H2 another core protein enhances the immunogenicity of A28, but is not itself a target of neutralizing antibody ^24, 35^. No information regarding immunogenicity was available for the other EFC proteins prior to this report. Our approach was to test sera of rabbits or mice that had been immunized with secreted ectodomains of EFC proteins for their abilities to neutralize VACV. We found that six EFC proteins are neutralization targets (L1, F9, A28, A16, G9 and J5) and four are non-neutralization targets (H2, G3, L5, and A21). O3, which is only 35 amino acids and composed mainly of a TM domain, was not tested. A16 and G9 induced high cross-neutralizing activities against VACV, CPXV and MPXV and were further analyzed. The antibodies prevented entry of VACV when added either before or after virus adsorption to cells. In a lethal mouse model, A16 and G9 provided protection against death and the A16/G9 heterodimer further provided protection against weight loss. The neutralizing activity induced by A16/G9 was greater than that elicited by MPXV M1, the homolog of VACV L1 that is a component of recent candidate mRNA vaccines and provided better protection than M1 in the mouse model. Furthermore, A16/G9 synergized with EV proteins providing protection as well or better than MVA. Taken together, A16 and G9 are good candidates for inclusion in new orthopoxvirus vaccines.

The neutralizing and non-neutralizing antibodies differ in their ability to bind intact virus particles. It was of interest to determine whether the locations of the proteins within the EFC could explain differences in antibody binding and neutralization. Although the 11 components of the EFC were identified by immunopreciptitation ^14^, isolation of the native complex has not been reported and the organization of the proteins has not been determined experimentally. For this reason we turned to predictive modeling initially using AlphaFold2 ^36^ and then AlphaFold3.

With AlphaFold2, we were able to model the 9 core proteins but a model for all eleven proteins was obtained here with AlphaFold3. The pLDDT and PAE metrics calculated by AlphaFold3 as well as the RMSD of the models compared to experimentally determined structures provided confidence in the core and holocomplex models. Importantly, the close associations of A16 and G9, A28 and H2, and L5 and G3 in the model is consistent with experimental data demonstrating that these pairs form stable heterodimers ^23, 30, 31^. Whether the model is most similar to a pre-or post-fusion structure is unknown. Nevertheless, the model provided a working template upon which we could superimpose neutralization and non-neutralization targets. The highly neutralizing antibody targets A16 and G9 formed the apex of the EFC, with the other targets of neutralizing antibodies just below. The four non-neutralizing targets were closely associated with each other and closer to the base of the complex suggesting that this region may be masked either by the other EFC proteins or by other viral membrane proteins. Nevertheless, virion-associated A21 and L5 (as well as F9, L1, A28 and G9) can react with the membrane-impermeable sulfo-NHS-SS-biotin ^20, 37–39^ indicating their accessibility by a small molecule.

Unexpectedly, antibodies binding to G9 and A16 were not detected in the blood of mice, rabbits and monkeys that had been vaccinated with attenuated or pathogenic VACV. These data are consistent with studies in which antibodies from humans that received the live virus smallpox vaccine were analyzed ^40^. Antibodies to the H3, A27, and L1 proteins were detected in 90.5%, 67.6 %, and 31% of vaccinees, respectively, whereas antibodies to A16 and G9 were found in none. Although subdominant protein regions that elicit only weak neutralizing antibody responses is common ^41^, the inability to detect binding antibodies to entire proteins displayed on the surface of viral particles seems exceptional.

One possibility to explain the poor immunogenicity of A16 and G9 during live virus infections is association with other immunodominant proteins in the viral membrane. Supporting this explanation are reports that, in contrast to other EFC proteins, A16 and G9 are not released from virus particles by incubation with DTT and the detergent NP40 ^39, 42^ and that A16/G9 stably interacts with viral proteins A56, K2, A25 and A26 ^30, 43^. Additionally, A16 and G9 can be chemically cross-linked to immunodominant H3, A25, D8, A26, and A27 in virus particles ^44^. To explore the idea that sequestering of A16 and G9 restricts their immunogenicity, we constructed a recombinant MVA in which G9 was expressed as a secreted protein and therefore not embedded in the viral membrane. In sharp contrast to the parental MVA, antibodies to G9 were detected after a single vaccination with the recombinant virus and were boosted after a second. As A16 and G9 are intrinsically immunogenic and antibodies to these proteins are highly neutralizing, the failure of live virus to induce antibodies to these proteins may represent a viral evasion mechanism. In this respect, recombinant vaccines that express A16 and G9 may have an advantage over live attenuated vaccines.

## Methods

### Biosafety

All work with VACV and CPXV and potentially infectious material derived from animals was performed at Biosafety level 2 and all work with MPXV was in a Biosafety level 3 select agent-approved laboratory at the NIAID, NIH in Bethesda, MD. Sample inactivation and removal was performed according to standard operating protocols approved by the local Institutional Biosafety Committee.

### Animal studies

All mouse experiments and procedures adhered to protocol LVD 12E approved by the NIAID Animal Care and Use Committee and followed NIH guidelines, the Animal Welfare Act, and U.S. federal law. Euthanasia was carried out at a 30% weight loss using carbon dioxide inhalation and cervical dislocation, in accordance with AVMA guidelines (2013 Report of the AVMA panel on euthanasia).

### AlphaFold prediction

Models were created with Alphafold version 3.0 using default settings with 5 random seeds per model. The highest ranking model was used for visualizations. The following protein sequences from the VACV WR strain (NC_006998) were used in the Alphafold predictions. A16:YP_233018.1, A21: YP_233022.1, A28: YP_233033.1, F9: YP_232930.1, G3:YP_232961.1, G9: YP_232969.1, H2: YP_232982.1, J5: YP_232979.1, L1: YP_232970.1, L5: YP_232974.1. Protein structures were visualized using Chimera X 1.9 https://pubmed.ncbi.nlm.nih.gov/37774136/. Superposition of the AlphaFold3 model with experimentally determined structures from PDB (A16:8GP6/A, A21:8U0R/A, A28:8GQO.1/A:30-109, F9:6CJ6/A, G3:7YTU/A, G9:8GP6/B, H2:8INI/A, J5:8WT5.1/A, L1:1YPY/A, L5:7YTU/B) was accomplished using Chimera X’s internal matchmaker command with default settings which carries out an iterative algorithm aligning C alpha atoms of matching residues and removing pairs with deviations larger than 2.0 Å resulting in a pruned alignment. Chimera X’s interfaces command was used to calculate the pairwise buried solvent-accessible areas (SASA) of all subunits in the complex. Again, default settings were used which resulted in a minimum interface size of 300 Å squared when measured with a 1.4 Å probe.

### Cells

African green monkey BS-C-1 cells (ATCC CCL-26) were cultured in modified Eagle minimal essential medium (EMEM) (Quality Biologicals, Inc., Gaithersburg, MD) supplemented with 10% fetal bovine serum (FBS) (Sigma-Aldrich, St. Louis, MO) at 37°C with 5% CO2 for virus replication and titration in Corning® sterile polycarbonate flasks equipped with 0.2 µm ventilated caps under 80% humidity. Adherent HEK293T cells were maintained in DMEM (Gibco) with 10% FBS at 37°C. Hela S3 cells (ATCC, CCL-2.2) were cultured in spinner-modified minimum essential medium (SMEM) with 1% Non-Essential Amino Acids, and 10% FBS at 37°C with 8% CO2. Both culture media were further supplemented with 2 mM L-glutamine, 100 U/mL penicillin, and 100 μg/ml streptomycin (Quality Biologicals, Inc.). Expi293F suspension cells (A14527, Thermo Fisher Scientific) were cultured in Expi293 medium (A1435101, ThermoFisher) at 37°C, 125 rpm, and a 5 to 8% CO2 atmosphere. All cell lines tested negative for mycoplasma.

### Viruses

VACV MVs were obtained by Dounce homogenization of HeLa S3 cells infected with VACV WRvFire ^45^, VACV-WR Gauss-A4 ^22^, CPXV-Brighton-GFP ^46^, and MPXV-Z-1979-GFP ^47^ at 48 h post-infection, following established protocols ^48^. MVA ^49^ and recombinant MVA-G9 were grown in CEF. MVs were purified by sedimentation through two 36% (w/v) sucrose cushions, followed by banding on a 25 to 40% (w/v) sucrose gradient ^48^. MVA and rMVA-G9 were titrated by plaque immunostaining in CEF cells ^48^. Purified stocks were stored at −80 °C and subjected to a 1-min sonication on ice prior to infection.

### Cloning, expression, and purification of recombinant proteins

To generate soluble VACV proteins, the C-terminal TM domains of A16 (residues 343-377; Uniprot accession number P16710 · A16_VACCW) and G9 (residues 320-340; Uniprot accession number P07611 · G9_VACCW) and the N-terminal TM domains of M5 (residues 52-128; Uniprot accession number PG099_MONPV) and G2 (residues 22-111; Uniprot accession number A0A0F6N9M0_MONPV) were excised. Codon-optimized sequences encompassing residues 3-342 of A16, residues 3-319 of G9, residues 52-128 of M5, residues 22-111 of G2 with modified N-glycosylation sites were synthesized and inserted into the pBudCE4.1 dual expression vector (ThermoFisher) downstream of the human modified albumin signal peptide ^50^.

Prior to transfection, Expi293F^TM^ cells were diluted to a density of 3 × 10^6^ viable cells/ml (95% viability) with fresh, pre-warmed Expi293^TM^ Expression Medium. For a 300 ml scale transfection, a total of 300 µg of plasmid was diluted with 18 ml Opti-MEMTM I Medium. Simultaneously, 960 µl ExpiFectamine^TM^ 293 Reagent was diluted with 16.8 ml Opti-MEM^TM^ I Medium. After a 5-min incubation at room temperature (RT), the Reagent was added to the plasmid DNA, mixed, and incubated at RT for an additional 15 min. The resulting ExpiFectamine^TM^ 293/plasmid DNA complexes were transferred to the cells. ExpiFectamine^TM^ 293 Transfection Enhancer 1 (1.8 ml) and 2 (18 ml) were added at 18–22 h post-transfection. The medium was harvested after 5 days and the proteins concentrated, passed through a 0.25 µm filter, and purified using gravity flow Ni-NTA affinity resin chromatography following the recommendations of QIAGEN. The proteins were eluted by overnight incubation at 4°C with 20 mM Tris/HCl, 200 mM NaCl, 150 mM imidazole; pH 7.5. The amount of purified proteins was determined using the Thermo Scientific Pierce BCA Protein Assay Kit.

### Immunoprecipitation

HEK293T cells expressing tagged proteins were suspended in ice-cold lysis buffer (ChromoTek), supplemented with a protease inhibitor cocktail and 1 mM PMSF. ChromoTek V5-Trap® Agarose, consisting of an anti-V5-tag nanobody (cat. no. v5ta, ChromoTek), or ChromoTek Flag-Trap® Agarose, consisting of an anti-flag-tag nanobody (cat. no. ffak, ChromoTek), were employed according to the manufacturer’s protocol for immunoprecipitation. The lysate was added to the equilibrated V5 or Flag beads and rotated end-over-end for 1 h at 4°C. After three washes, the beads were resuspended in 2x SDS-sample buffer and used for immunoblot analysis.

### Immunoblot of proteins

Proteins were resolved by SDS-PAGE and transferred to nitrocellulose membranes. The membranes were blocked at RT for 1 h in phosphate-buffered saline (PBS) with Tween 20 (PBS-T), supplemented with 5% nonfat dry milk and then incubated at 4°C overnight in PBS-T with primary antibodies directed against V5 (cat. no. ab184331, 1:1000, Abcam), Flag (cat. no. F1804, 1:5,000, Sigma-Aldrich), His (Proteintech, cat. no. 66005-1-Ig, 1:10,000), GAPDH (cat. no. 60004-1-Ig, 1:20,000, Proteintech), actin (cat. no. 60004-1-Ig, 1:20,000, Proteintech). After three washes, the membranes were incubated with IRDye 680RD goat anti-mouse or rabbit IgG secondary antibody (cat. no. 926-68070, 1:10,000, Li-Cor Biosciences) or IRDye 800CW goat anti-rabbit or mouse IgG secondary antibody (cat. no. 926-32211, 1:10,000, Li-Cor Biosciences), or goat-anti-mouse or anti-rabbit horse raddish peroxidase (HRP) secondary antibody. The secondary anti-mouse antibody was applied at a dilution of 1:2000, followed by detection utilizing an Odyssey infrared scanner (Li-Cor Biosciences). When secondary antibodies with HRP or only anti-flag HRP were used, the chemiluminescence substrate (Radiance ECL, azure biosystems) was applied to the membranes, and the immunoblot was subsequently resolved using the Li-Cor Biosciences or Azure biosystems Imager system.

### Depletion of specific antibodies from serum

100 µl (4 mg) of TALON Dynabeads (Invitrogen) were incubated with 100 µg of purified histidine-tagged A16, G9, or A16/G9 proteins for 4 h at 4°C. The beads were washed and subsequently incubated overnight with 500 µl of 1:200 diluted A16/G9 serum in PBS at 4°C. Following incubation, the beads were removed from the serum magnetically. The protein-adsorbed sera were then utilized for antibody binding and neutralization assays.

### Immunization and challenge of mice

Female 6 to 7 weeks BALB/c mice from Taconic Biosciences were caged in groups of five. The mice were maintained in small, ventilated microisolator cages within an animal BSL-2 pathogen-free environment. Male mice were not used because of their aggressive behavior, which would require housing in additional cages that were not available. Proteins (10 μg per mouse) were diluted in 50 μl of PBS and emulsified with an equal volume of AddaVax adjuvant (InvivoGen). Subcutaneous injections were administered at 3-week intervals. For live virus immunization, 10^7^ PFU of MVA in PBS supplemented with 0.05% BSA were injected IM into each hind leg of the mouse. Mice were challenged IN with 10^5^ PFU or 10^6^ VACV of WRvFire after the third immunization. Daily weight measurements were recorded, and mice were euthanized if they lost 30% of their initial body weight.

### Bioluminescence imaging

Mice were anesthetized with isoflurane and injected IP with Xenolight d-luciferin substrate (150 μg/g body weight) 8 to 10 min before imaging with an IVIS Lumina LT series III system (Perkin Elmer, Waltham, MA) as described ^28^. Luminescent exposures (1-60 sec) were collected using small or medium binning factors and diverse f-stop settings. Living Image Software (Perkin Elmer) facilitated acquisition and analysis. Photon flux was measured by outlining the head and body separately as regions of interest, quantifying light emission in photons per second per square centimeter per steradian.

### ELISA

VACV F9 (BEI Resources: NR-2626), MPXV H3 (Sino Biological, 40893-V08H1) and MPXV M1 (Sino Biological, 40904-V07H) were diluted in 0.05 M carbonate-bicarbonate buffer, pH 9.6, to achieve a concentration of 1 µg/ml. A high-binding 96-well ELISA plate (Clear Round-Bottom Immuno Nonsterile 96-Well Plates, 3655TS) was filled with 100 µl of the diluted protein (0.1 µg/well) and incubated overnight at 4 °C. Following adsorption, the wells were washed thrice with 1X wash buffer (prepared from 10X wash buffer: 270 g NaCl and 30 mlTween-20 in 3l H_2_O). Subsequently, the plates were blocked for 2 h at room temperature with 100 µl of 0.2 ml Tween-20 and 5 g non-fat dry milk in 100 ml PBS and washed three times with 1X wash buffer. A 96-well ELISA plate was coated with 60 μl/well (120 ng/well) of streptavidin solution at a concentration of 2 μg/ml in PBS, incubated at 4°C overnight, and washed twice with 1X wash buffer. Wells were incubated with 100 μl/well of block buffer for 1 h at RT. Biotinylated A16, G9, and A16/G9 dimers (0.5 μg/ml in block buffer), and A28 and H2 (0.25 μg/ml in block buffer), were added to the wells (100 μl/well or 25 ng/well) for 45 min The plate was washed four times with 1X wash buffer. Sera from vaccinated mice were heat-treated at 56 °C for 30 min to inactivate complement, and a series of eight 2-fold dilutions of mouse sera were added, with incubation continued for 1 h at RT. Plates were then incubated with goat anti-mouse IgG-horseradish peroxidase antibody in a 1:2,000 dilution (Thermo Fisher) for 1 h at RT and developed with 100 µl of pre-warmed SureBlue TMB substrate (SeraCare) for 30 min at RT after washing three times with 1X wash buffer. Absorbance was measured at 370 and 492 nm using a Synergy H1 plate reader with Gen5 analysis software (Agilent Technologies). EC50 values were determined using Prism.

For intact virus ELISA, sucrose gradient-purified VACV WR was added to plates at a final concentration of 2 × 10^7^ pfu/ml (100 µl) in 0.05 M carbonate-bicarbonate buffer, pH 9.6, and incubated overnight at 4 °C. Unbound virus was removed from plates and discarded in chlorox, followed by the addition of 100 µl of 2% paraformaldehyde to each well to inactivate bound virus. Plates were incubated at 4°C for 10 min, washed five times with 1X wash buffer, and rinsed with dH_2_O. Then, 200 µl of block buffer was added to each well and incubated at 37°C for 1 h. The procedure for adding diluted serum was the same as previously described above.

For dissociated virus ELISA, purified VACV particles (10^9^ PFU/ml) were incubated in PBS with or without 1% SDS for 1 h at room temperature. The samples were then diluted 50-fold in 0.05 M carbonate-bicarbonate buffer (pH 9.6) supplemented with 0.5% BSA and aliquots (100 µl) were added to plates and incubated overnight at 4 °C. Unbound virus was removed and discarded in bleach. 100 µl of 2% paraformaldehyde was added to each well, followed by incubation at 4 °C for 10 min. Plates were washed five times with 1× wash buffer and rinsed with distilled water. Wells were then blocked with 200 µl of blocking buffer for 6 h at 4 °C.

### Neutralization Assay

A 96-well plate semi-automated flow cytometric assay was used to quantitatively assess VACV-WR and MPXV Z-1979 expressing Aequorea coerulescens GFP, following established procedures ^21^. Two-fold dilutions of heat-inactivated immune serum from individual mice were prepared in 96-well, round-bottom polypropylene plates using SMEM containing 2% FBS.

Approximately 2.5 × 10^4^ PFU of VACV-WR or MPXV-Z-1979 or CPXV-Brighton GFP-expressing viruses were added to each well, followed by a 1-h incubation at 37°C. Afterward, 10^5^ HeLa S3 cells were added to each well, and plates were incubated for an additional 16 to 18 h at 37°C. Fixation of the cells with 2% paraformaldehyde inactivated the virus, and GFP expression in infected cells was quantified using a fluorescence-activated cell sorting Canto II flow cytometer, and the acquired data were analyzed using FlowJo software (BD Biosciences). NT_50_ values, representing the serum dilution at which 50% neutralization occurred, were determined using Prism software (GraphPad/Dotmatics).

### Cell binding inhibition-Gaussia luciferase flash assay

HeLa S3 cells (1.25 × 10^5^) were seeded in a 96-well plate. VACV Gaussia-A4 virions (5 PFU per cell) were pre-treated with: control PBS-mock immunized mouse serum, A16, G9, A16/G9 and VACV immune mouse sera (1:100 dilution). The virions were then incubated with the cells at neutral pH for 1 h at 4°C. After incubation, the cells were washed twice with cold PBS to remove unbound virus and then lysed using 1X Cell Lysis Buffer (Thermo Scientific, Pierce™ Gaussia Luciferase Flash Assay Kit) for 15 min at RT. Gaussia luciferase activity was measured using 10 μl/well of cell lysate and 50 μl 1X Coelenterazine in a Berthold Sirius luminometer.

### Pre-and post-attachment entry inhibition assays

Pre-attachment inhibition assay: 50 µl of diluted serum from immunized mice was incubated with 1.25×10^5^ PFU of purified WRvFire at a multiplicity of infection (MOI) of 1 in EMEM + 2.5% FBS at room temperature for 30 min. Following this, the mixtures were added to pre-chilled Hela S3 cells (1.25×10^5^ cells per well) in cold EMEM + 2.5% FBS at 4°C for 1 h to allow attachment of the virus or virus-antibody complex to the cell surface. The cells were washed twice with cold PBS by centrifuging at 2,500 rpm for 5 min at 4°C to remove unbound virions. Subsequently, the cells were incubated with pre-warmed EMEM + 2.5% FBS for 2 h at 37°C. After this incubation, the cells were washed with PBS at 2,500 rpm for 5 min and treated with 100 µl of 1X Cell Culture Lysis Reagent (Promega) for 20 min at RT with gentle agitation. Following centrifugation at 12,000 rpm for 10 min, 20 µl of the lysate was mixed with 100 µl of substrate. Luciferase activity in cellular extracts was measured according to the Promega protocol and quantified using a Berthold Sirius luminometer (Berthold Detection Systems).

Post-attachment inhibition assay: 1.25×10^5^ Hela S3 cells were seeded in 96-well plates. Purified WRvFire (3.75×10^5^ PFU, MOI=3) were incubated with Hela S3 cells at 4°C for 30 min to allow attachment. Subsequently, the cells washed twice with cold PBS to eliminate unbound virus and then were incubated with 50 μl of serially diluted immune or control mouse serum for 30 min at 4°C. After an additional two washes with cold PBS, the cells were incubated at 37°C for 2 h and luciferase activity determined.

### Construction of recombinant MVA

The recombinant MVA expressing G9 (rMVA-G9) was constructed by homologous recombination. The G9R ORF was inserted into the pLW44 vector under the control of the synthetic promoter mH5, the GFP reporter gene was regulated by the p11 promoter in the pLW44 vector for selection ^51^. CEF were infected with MVA at an MOI of 2 for 2 h at 37°C and transfected with pLW44-G9R followed by 2-days incubation at 37°C with 5% CO2. Integration of the GFP reporter into the MVA genome was confirmed by fluorescence using an EVOS fluorescence microscope. To eliminate wild-type MVA, viral seeds underwent repeated plaque purification. Clones from the fifth round of purification were selected and expanded for subsequent purification and assessment of G9 expression. Validation procedures included observation of GFP fluorescence and detection of Avi-tagged G9 secretion via immunoblot analysis in infected cell supernatants. Confirmation of G9 insertion within the rMVA-G9 genome was achieved through viral DNA extraction and PCR. After amplification, rMVA-G9 was sucrose gradient purified.

## Data analysis

Data were analyzed with GraphPad Prism 10 software (CA, USA) using one-way ordinary ANOVA with multiple comparisons.

## Data availability

All data and unique materials supporting the findings of this study are available within the paper or from the corresponding author upon request.

## Supporting information

Movie 2. EFC holocomplex showing neutralizing targets in color and non-neutralizing regions in gray.

## Acknowledgements

Support was provided by the Division of Intramural Research, NIAID, NIH. The AlphaFold3 model was generated on the computational resources of the NIH HPC Biowulf cluster (https://hpc.nih.gov).

## Author contributions

HY and BM designed the experiments; HY, CAC, WX, MAI, PLE and TK performed experiments; WR carried out the protein prediction analysis; GHC and AAB provided materials; HY and BM wrote the manuscript;. BM supervised the project. All authors discussed the results and commented on the manuscript.

## Competing interests

HY, BM and WR are inventors on US government patent applications.

## Figure Legends

**Fig. S1.**
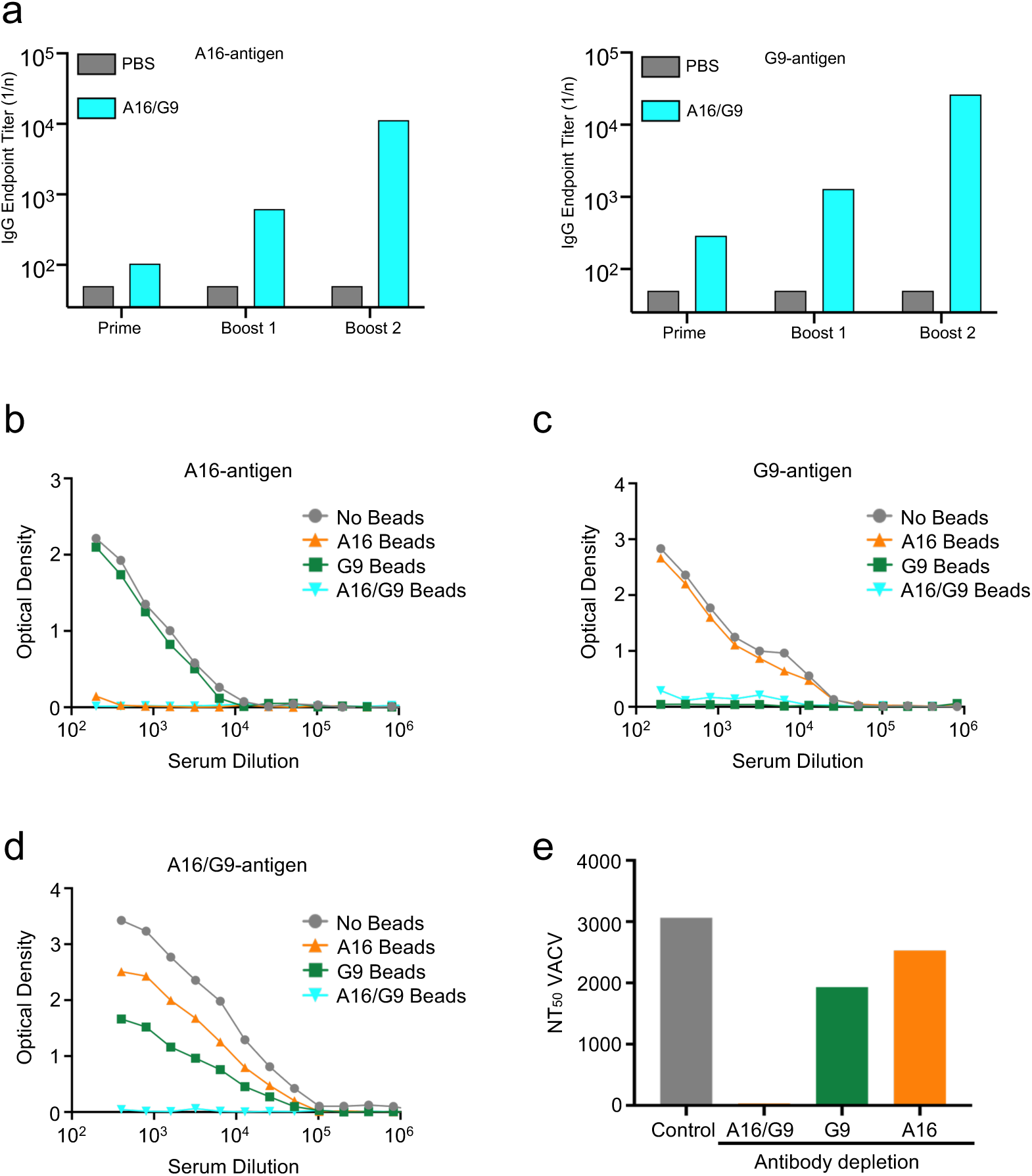
Binding and depletion of antibodies from mice immunized with A16/G9. **(a)** Binding of A16/G9 serum to A16 and G9 antigens. **(b-d)** Depletion of A16, G9 and A16/G9 by proteins bound to NiNTA beads determined by ELISA. **(e)** Depletion of VACV neutralizing activity in serum of mice immunized with A16/G9 heterodimers.

**Fig. S2.**
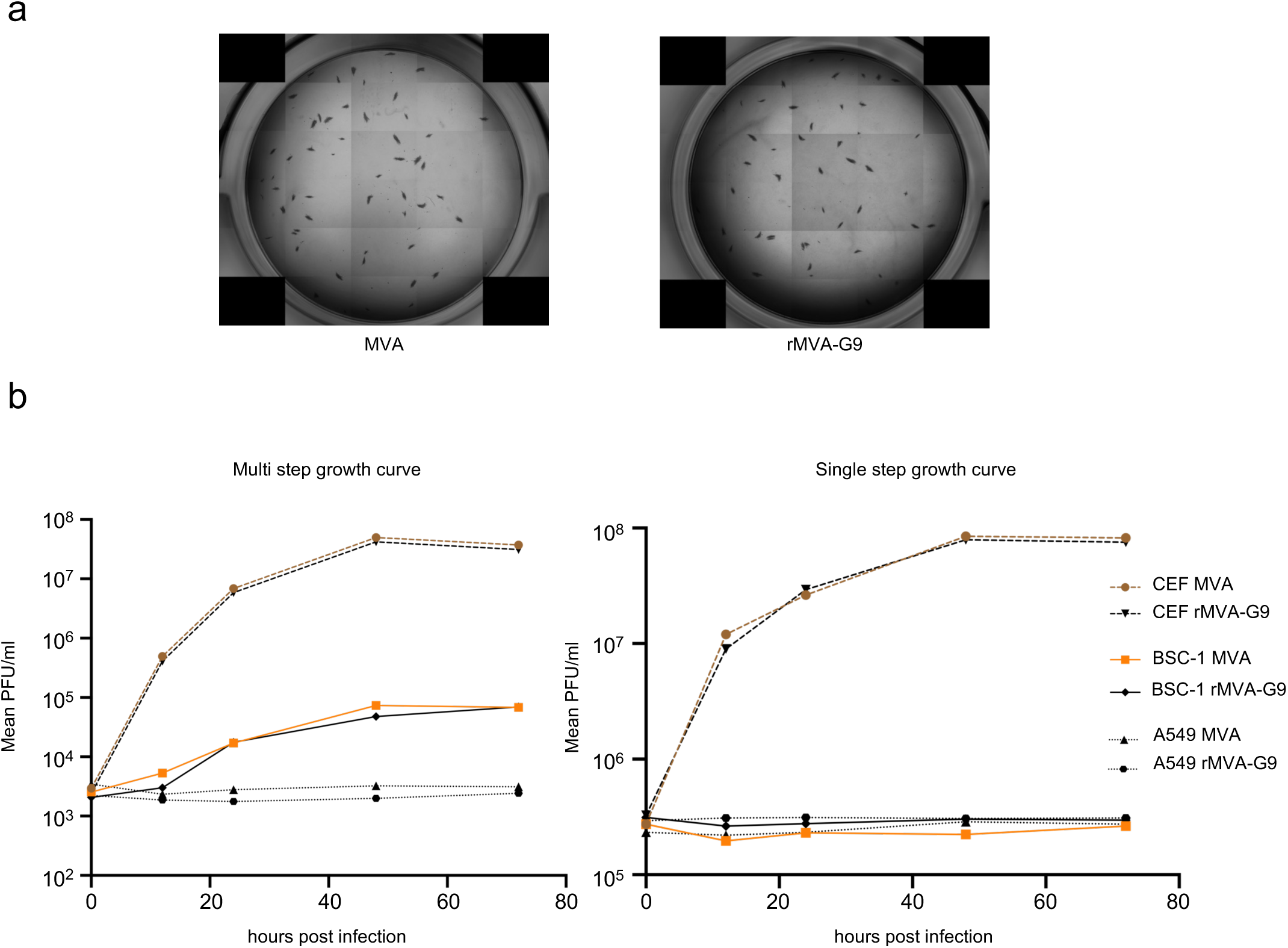
Replication of recombinant MVA expressing secreted G9. **(a)** Virus foci formed in CEF monolayers within 48 h and stained with anti-VACV antibodies. **(b)** Multistep (0.1 PFU) and single step (10 PFU) growth curves of MVA and recombinant MVA following infection of BS-C-1, CEF and A549 cells.

**Fig. S3.**
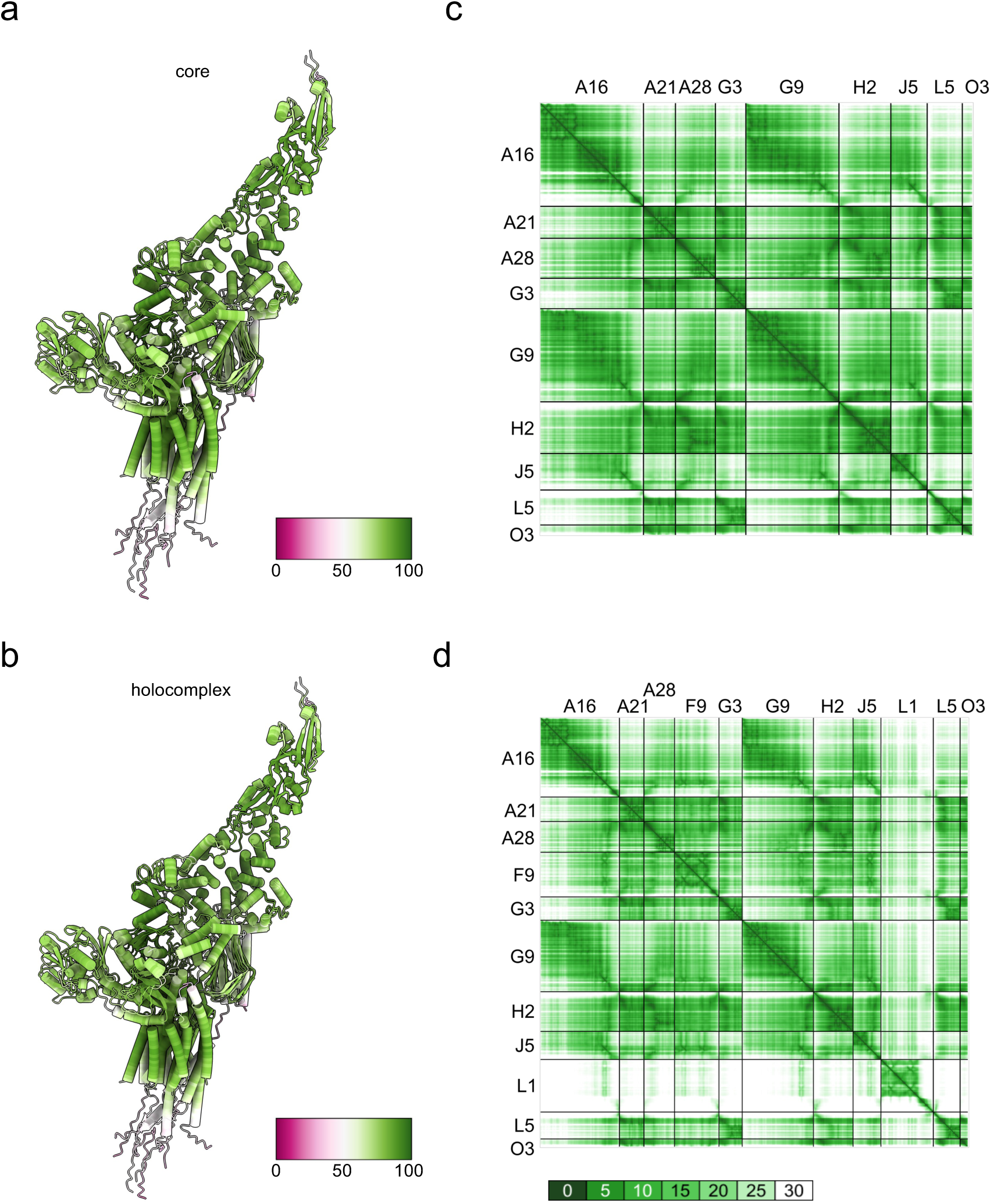
Confidence metrics of predicted core and holocomplex EFC models. The predicted local difference test (pLDDT) per-residue measure of local confidence of core **(a)** and holocomplex **(b)** models scaled 0 to 100 with higher score indicating higher confidence. Predicted aligned error (PAE) measure of confidence in relative position of two residues within predicted core **(c)** and holocomplex **(d)** models. Regions of high confidence are dark green.

**Fig. S4.**
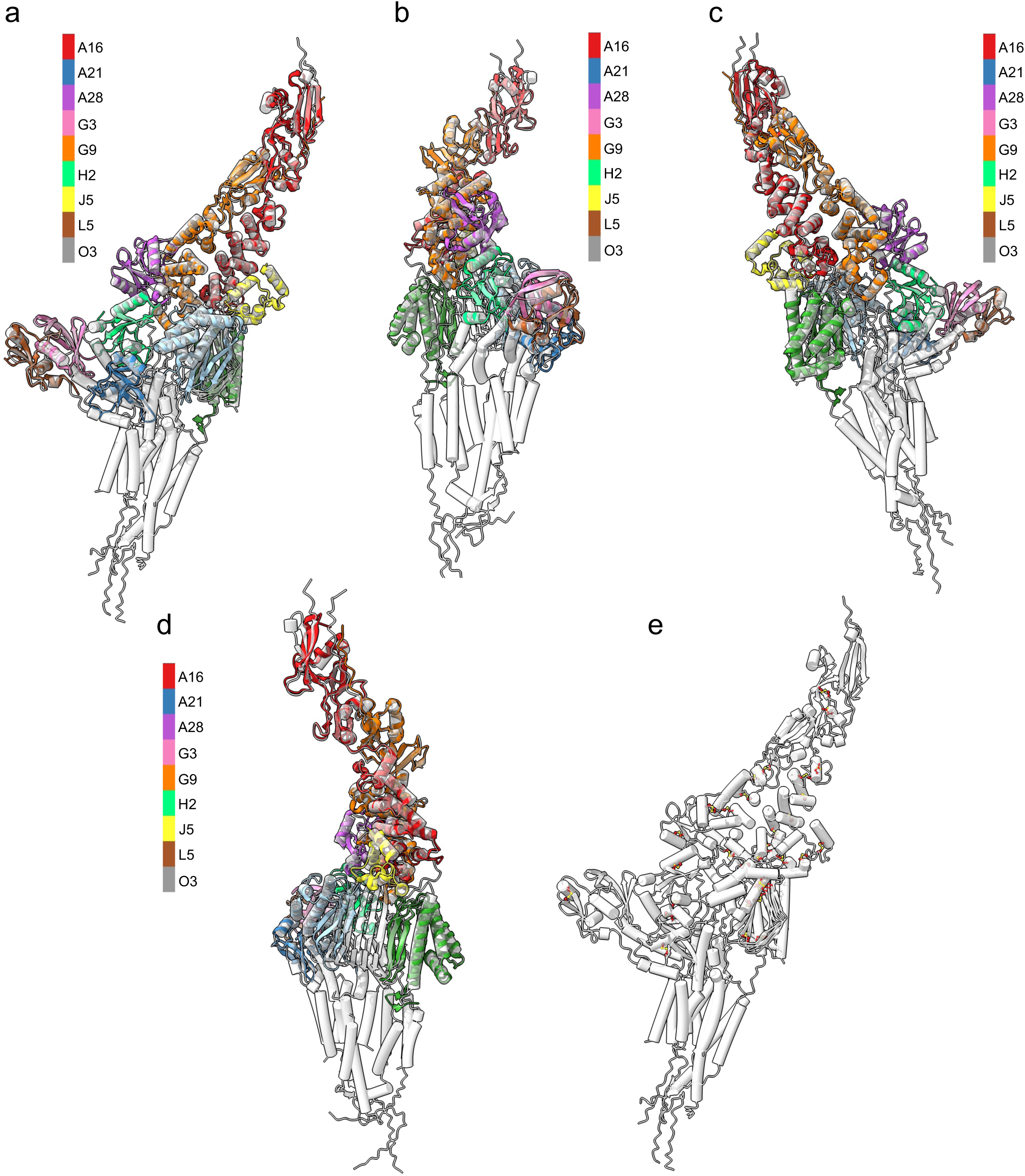
Correlation of predicted and experimental structures. (**a-d**) The predicted holocomplex in light gray with superimposed ectodomain structures from the Protein Database (PDI) is shown in four 90° rotations. Individual proteins identified by color. **(e)** Close association of cysteine residues in holocomplex model. Pairs of cysteines in red.

**Fig. S5.**
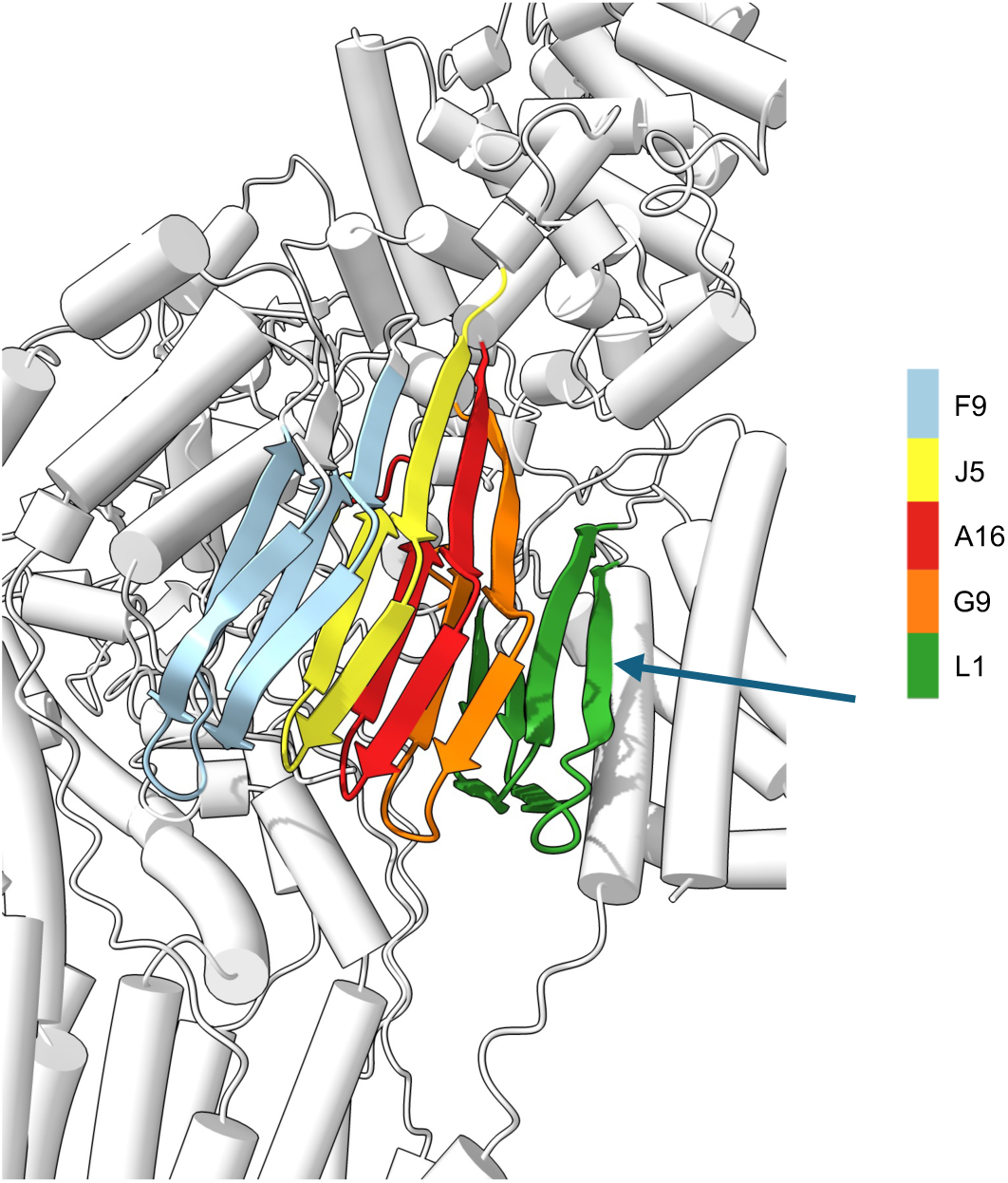
Model of holocomplex showing alignment of β-folds of A16, G9, J5 and F9. β-folds in color.

**S1 movie**. **EFC holocomplex**. 3-dimensional space-filling model of the holocomplex with surface lipophilicity, and parallel β-sheets shown.

**S2 movie. EFC holocomplex showing neutralizing targets in color and non-neutralizing regions in gray**.

## Notes

### Competing Interest Statement

HY, WR, BM are inventors on vaccine applications by US government

